# Understanding sequencing data as compositions: an outlook and review

**DOI:** 10.1101/206425

**Authors:** Thomas P. Quinn, Ionas Erb, Mark F. Richardson, Tamsyn M. Crowley

## Abstract

**Motivation:** Although seldom acknowledged explicitly, count data generated by sequencing platforms exist as compositions for which the abundance of each component (e.g., gene or transcript) is only coherently interpretable relative to other components within that sample. This property arises from the assay technology itself, whereby the number of counts recorded for each sample is constrained by an arbitrary total sum (i.e., library size). Consequently, sequencing data, as compositional data, exist in a non-Euclidean space that renders invalid many conventional analyses, including distance measures, correlation coefficients, and multivariate statistical models.

**Results:** The purpose of this review is to summarize the principles of compositional data analysis (CoDA), provide evidence for why sequencing data are compositional, discuss compositionally valid methods available for analyzing sequencing data, and highlight future directions with regard to this field of study.

## 1 From raw sequences to counts

Automated Sanger sequencing served as the primary sequencing tool for decades, ushering in significant accomplishments including the sequencing of the entire human genome ([50]). Since the mid-2000s, however, attention has shifted away this “first-generation technology” toward new technologies collectively know as next-generation sequencing (NGS) ([50]). A number of NGS products exist, each differing in the sample preparation required and chemistry used ([50]). Although each product tends toward a different application, they all work by determining the base order (i.e., sequence) from a population of nucleotides, such that it becomes possible to estimate the abundances of unique sequences ([50]). However, these sequence abundances are not absolute abundances because the total number of sequences measured by NGS technology (i.e., the library size) ultimately depends on the chemistry of the assay, not the input material.

Depending on the input material, NGS has many uses. These include (1) variant discovery, (2) genome assembly, (3) transcriptome assembly, (4) epigenetic and chromatin profiling (e.g., ChIP-seq, methyl-seq, and DNase-seq), (5) meta-genomic species classification or gene discovery, and (6) transcript abundance quantification ([50]). The application of NGS to catalog transcript abundance is better known as RNA-Seq ([50]) and can be used to estimate the portional presence of transcript isoforms, gene archetypes, or other. RNA-Seq works by taking a population of (total or fractionated) RNA, converting them to a library of cDNA fragments, optionally amplifying the fragments, and then sequencing those fragments in a “high-throughput manner” ([73]). When sequencing smaller RNA (e.g., microRNA), an additional size selection step is used to ensure a uniform size of the RNA product ([36]).

The result of RNA-Seq is a virtual “library” of many short sequence fragments that are converted to a numeric data set through alignment (most often to a previously established reference genome or transcriptome) and quantification ([33]). The alignment and quantification steps summarize the raw sequence data (i.e., reads) as a “count matrix”, a table containing the estimated number of times a sequence successfully aligns to a given reference annotation. The “count matrix” therefore provides a numeric distillation of the raw sequence reads collected by the assay; as such, it constitutes the data routinely used in statistical modeling, including differential expression analysis ([33]). Two factors complicate alignment and quantification. First, assembled references (e.g., genomes or transcriptomes) are only just references: sequences measured from biological samples will have an expected amount of variation, either systematic or random, when compared with the reference. This variation necessitates that the alignment procedure accommodates (at least optionally) a certain amount of mismatch ([16]). Meanwhile, some reads (notably short reads) can ambiguously map to multiple reference sites, an undesired outcome that is amplified by mismatch tolerance ([16]). Many alignment and quantification methods exist (e.g., TopHat ([68]), STAR ([18]), Salmon ([51]), and others) and are reviewed elsewhere (e.g.,[29]; [27]; [42]; [21]; [34]; [9]; [72]; [8]).

The “count matrix” (or equivalent) produced by alignment and quantification is routinely analyzed using statistical hypothesis testing (e.g., generalized linear models) or data science techniques (e.g., clustering or classification). Most commonly, data are studied using differential expression analysis, a constellation of methods that seek to identify which unique sequence fragments (if any) differ in abundance across the experimental condition(s). Like alignment and quantification, many differential expression methods exist (e.g., Cufflinks ([69]), limma ([55]), edgeR ([56]), DESeq ([7]), and others) and are reviewed elsewhere (e.g., [17]; [54]; [63]; [26]; [35]; [59]; [61]; [65]; [49]). However, it is important to note that conclusions drawn from RNA-Seq data appear to have a certain “robustness” to the choice in the alignment and quantification method, such that the choice in the differential expression method impacts the final result most ([75]).

The focus of this review is not to elaborate on the niceties of alignment, quantification, or differential expression, but rather to discuss the relative (i.e., compositional) nature of sequencing count data and the implications this has on many analyses (including differential expression analysis). In this review, we show how sequencing count data measure abundances as portions, rendering many conventional methods invalid. We then discuss methods available for dealing with portional data. Finally, we conclude by discussing challenges specific to these analyses and by considering advancements to this field of study. Although we emphasize RNA-Seq data throughout this paper, the principles discussed here apply to any NGS abundance data set.

## 2 Counts as parts of a whole

### 2.1 Image brightness as portions

As an analogy, let us imagine that we instructed two photographers to take a series of black and white photographs using a digital camera. We can represent the captured images as a set of *N*-dimensional vectors where each element (i.e., pixel) records the amount of light that hit a corresponding part of the film sensor. Considering this data set, let us ask a pointed experimental question: which photographer captured their photographs in brighter light? Better yet, for which pixels, on average, did Photographer A capture brighter light than Photographer B?

On first glance, this appears straight-forward. However, we want to know about the amount of light present when the photograph was taken, not the amount of light recorded by the film sensor. Although related, many factors influence the light measured at a given pixel. These include, for example, exposure time, aperture diameter, and the sensitivity of the film sensor. Changing any one of these parameters will change the image. Of course, such a change in the image does not mean a change in the reality.

At each pixel, we could then define two variables: luminance, the amount of light present at the moment of the photograph, and brightness, the amount of light perceived by the film sensor. Intuitively, we can understand brightness (the observed value, *o*), as a function, *f*, of luminance (the actual value, *a*:)

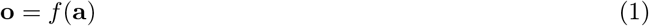

Even if we do not know the function, *f*, that relates these two measures, we see here that the total brightness recorded (i.e., Σ o) is an artifact of the conditions under which the luminance is measured. Yet, if we can assume that the film sensor responds proportionally to light and does not clip (an unrealistic and idealized assumption), then the portional brightness would equal the portional luminance:

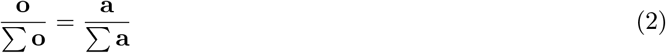

In this scenario, we can understand each element of o as a portion of the whole. As such, the brightness of a single pixel is only meaningful when interpreted relative to the total brightness (or to the brightness of the other pixels). Importantly, it follows that the ratio of any two parts of brightness will equal the ratio of any two parts of luminance.

### 2.2 Sequence abundance as portions

RNA-Seq data, through alignment and quantification, measure transcript abundance as counts. However, like the brightness of a digitalized image, the amount of RNA estimated for each transcript depends on some factors other than the amount of RNA molecules present in the assayed cell. Like a photograph, it is possible to change the observed magnitude while keeping the actual input the same. As such, RNA-Seq count data are not actually counts *per se,* but rather portions of a whole.

**Figure 1:**
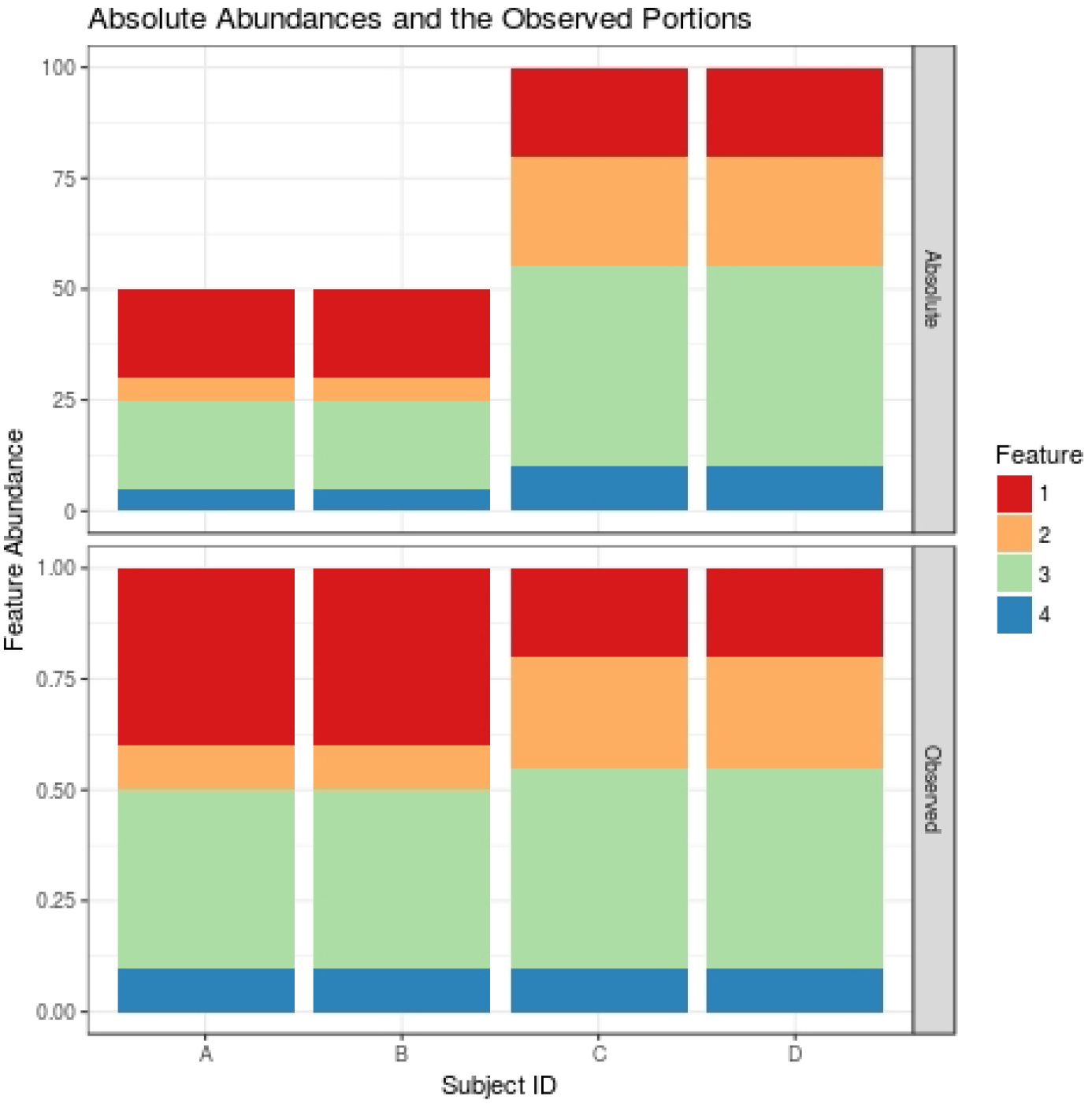
The top panel shows the absolute abundances of 4 features measured across 4 samples. The bottom panel shows their relative (i.e., portional) abundances for those same samples. Although Feature 4 is less abundant in samples A and B, it appears equally abundant across all samples when viewed as portions. Although Feature 1 is equally abundant across all samples, it appears more abundant in samples A and B when viewed as portions.

In fact, this is a property of all NGS abundance data: the abundances for each sample are constrained by an arbitrary total sum (i.e., the library size) ([63]). Since the library size is arbitrary, the individual values of the observed counts are irrelevant. However, the relative abundances of the observed counts still carry meaning. We can understand this by considering how, for a given sample, o, the library size (i.e., Σ o) cancels for a ratio of any two transcripts, *i* and *j*:

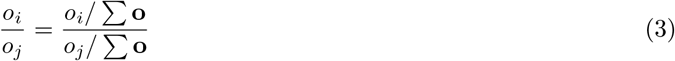

Analogous to how the relationship between luminance and brightness is unique to each photograph, the relationship between the actual abundances and the observed abundances is unique to each sample. Each independent sample, whether derived from a human subject or a cell line, may have undergone systematic or random differences in processing at any stage of RNA extraction, library preparation, or sequencing, causing between-sample biases ([63]). As such, library sizes typically differ between samples, making direct comparisons impossible ([63]). However, because the counts are portions of a whole, the interpretation is complicated even when library sizes are constant. For example, a large increase (or large decrease) in only a few transcripts will necessarily lead to a decrease (or increase) in all other measured counts ([63]). Figure 1 provides an abstracted visualization of how this might happen.

## 3 Counts as compositional data

### 3.1 The definition of compositional data

Compositional data measure each sample as a *composition,* a vector of non-zero positive values (i.e., components) carrying relative information ([2]). Compositional data have two unique properties. First, the total sum of all component values (i.e., the library size) is an artifact of the sampling procedure ([71]). Second, the difference between component values is only meaningful proportionally (e.g., the difference between 100 and 200 counts carries the same information as the difference between 1000 and 2000 counts ([71]).

Examples of compositional data include anything measured as a percent or proportion. It also includes other data that are incidentally constrained to an arbitrary sum. NGS abundance data have compositional properties, but differ slightly from the formally defined compositional data in that they contain integer values only. However, except for possibly at near-zero values, we can treat so-called *count compositional data* as compositional data ([43]; [53]). Note that it is not a requirement for the arbitrary sum to represent complete unity: many data sets (including possibly NGS abundance data) lack information about potential components and hence exist as incomplete compositions ([1]).

### 3.2 The consequences of compositional data

Compositional data do not exist in real Euclidean space, but rather in a sub-space known as the simplex ([2]). Yet, many commonly used metrics implicitly assume otherwise; such metrics are invalid for relative data. This includes distance measures, correlation coefficients, and multivariate statistical models ([12]). For compositional data, the distance between any two variables is erratically sensitive to the presence or absence of other components ([4]). Meanwhile, correlation reveals spurious (i.e., falsely positive) associations between unrelated variables ([52]). In addition, multivariate statistics yield erroneous results because representing variables as portions of the whole makes them mutually-dependent, multivariate objects (i.e., increasing the abundance of one decreases the portional abundance of the others) ([12]). All of this applies to NGS abundance data too ([43]).

In the life sciences, count data are usually modeled using the Poisson distribution or negative binomial distribution ([11]). For NGS abundance data, the negative binomial model is preferred because it accommodates situations in which the variance is much larger than the mean, a common feature of biological replicates in RNA-Seq studies ([63]). These models are necessary because analyzing non-normalized and non-transformed count data as if they were normally distributed would imply that it is possible to sample negative and non-integer values, contradicting the assumptions behind many statistical hypotheses ([15]) (although it is possible to extend Gaussian analysis to counts by use of precision weights ([39])). Moreover, NGS abundance data are compositional counts, not counts, meaning that the measured variables (i.e., components) are not univariate objects ([13]).

### 3.3 Normalization to effective library size

Although the negative binomial distribution is still used to model NGS abundance data ([63]), doing so necessitates (at the very least) an additional normalization step ([63]). The simplest normalization would involve rescaling counts by the library size (i.e., the total number of mapped reads from a sample) ([63]), but this does not transform compositional counts into absolute counts. Instead, analysts most often use other, more elaborate normalization methods that (generally speaking) adjust the individual counts of each sample based on the counts of a reference (or pseudo-reference) sample ([17]). The sum of these rescaled counts is called the effective library size.

Effective library size normalization for RNA-seq data was first proposed in an attempt to address the relative (i.e., closed) nature of the data through a method known as the trimmed mean of M (TMM) ([57]). This normalization works by inferring an ideal (i.e., unchanged) reference from a subset of transcripts based on the assumption that the majority of transcripts remain unchanged across conditions. Here, the reference was chosen to be a trimmed mean ([57]), although others have proposed using the median over the transcripts as the reference ([7]). The TMM normalizes data to an effective library size based on the principle that if counts are evaluated relative to (i.e., divided by) an unchanged reference, the original scale of the data is recovered. In the language of compositional data analysis, this approach is described as an attempt to “open” the closed data, and is often criticized on the basis that “there is no magic powder that can be sprinkled on closed data to make them open” ([3]). Yet, if the data were open originally (and only incidentally closed by the sequencing procedure), this point of view is perhaps extreme. On the other hand, if the cells themselves produce closed data by default (e.g., due to their limited capacity for mRNA production ([60])), any attempt to open the data might prove futile.

Given the difficulties in identifying a truly unchanged reference (and in interpreting it correctly in the case that closed data is being produced by the cells themselves) avoiding normalization altogether would seem desirable. After all, the choice of normalization method impacts the final results of an analysis. For example, the number and identity of genes reported as differentially expressed change with the normalization method ([41]), as do false discovery rates ([40]). This also holds true for compositional metabolomic data ([58]). Moreover, at least some normalization methods are sensitive to the removal of lowly abundant counts ([41]), as well as to data asymmetry ([63]).

**Figure 2:**
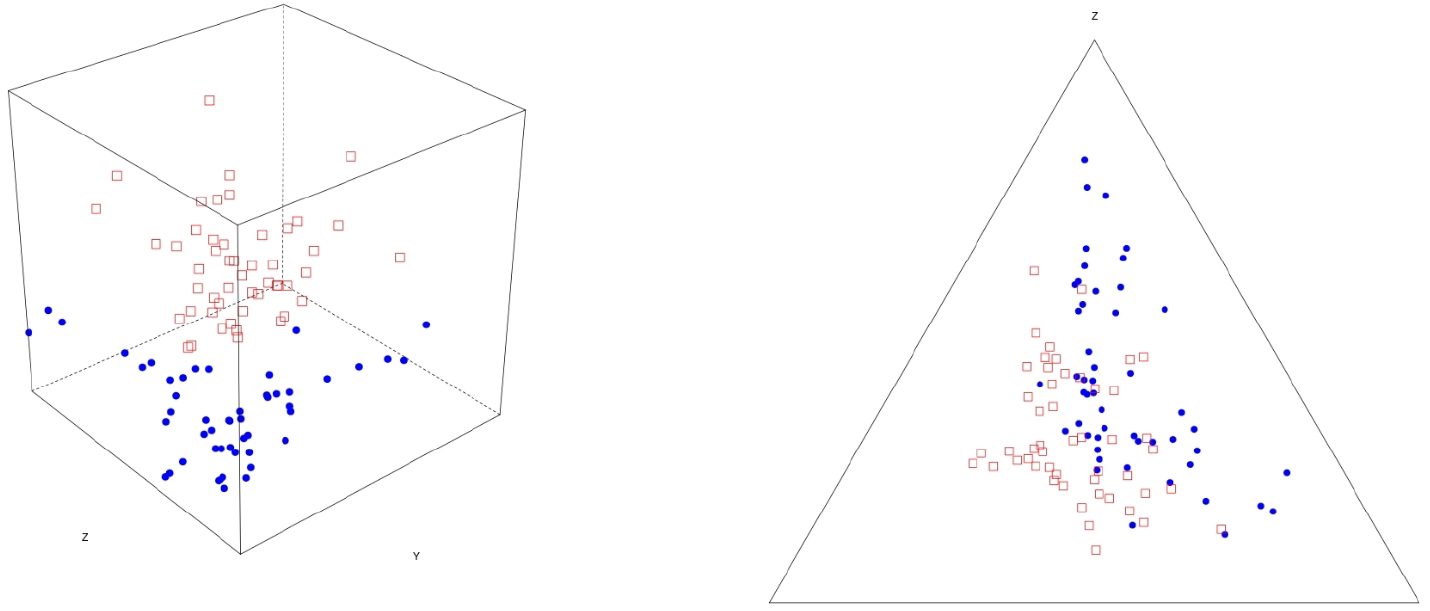
The left panel shows a 3D scatter of 100 samples, belonging to one of two groups, measuring the absolute abundances of 3 features. The right panel shows a ternary diagram of those same samples with the
3 features measuring relative (i.e., portional) abundances. Although the difference between the two groups is apparent in absolute space, the boundary between them becomes unclear in relative space. Note that for these relative data, it is not possible to reclaim a clear separation of the groups through transformation owing to the limited number of features available and the magnitude of the noise.

## 4 Principles of compositional data analysis

### 4.1 Approaches to compositional data

In lieu of normalization, many compositional data analyses begin with a transformation. Although compositional data exist in the simplex, Aitchison first documented that these data could get mapped into real space by use of the log-ratio transformation ([2]). By transforming data into real space, measurements like Euclidean distance become meaningful ([4]). However, it is also possible to analyze compositional data without log-ratio transformations. One approach involves performing calculations on the components themselves (called the “staying-in-the-simplex” approach) ([47]). Another involves performing calculations on ratios of the components (called the “pragmatic” approach) ([32]). Nevertheless, many compositional data analyses still begin with a log-ratio transformation.

Unlike normalizations, log-ratio transformations do not claim to open the data. Instead, the interpretation of the transformed data (and some of their results) depend on the reference used. In contrast, normalizations assume that an unchanged reference is available to recover the data (i.e., up to a proportionality constant) as they existed prior to closure by sequencing. Yet, while log-ratio transformations are conceptually distinct from normalizations, they are sometimes interpreted as if they were normalizations themselves ([25]). Although this contradicts compositional data analysis principles, conceiving of transformations as normalizations is helpful in understanding their use in some RNA-Seq analyses. Such log-ratio “normalizations”, like conventional normalizations, aim to recast compositional data in absolute terms, allowing for a straight-forward univariate interpretation of the data. Like effective library size normalization, this is done through use of an ideal reference.

### 4.2 The log-ratio transformation

First, let us consider a small relative data set with only 3 features measured across 100 samples. These samples belong to one of two groups. One of the features, “X”, can differentiate these groups perfectly. The other features, “Y” and “Z”, constitute noise. We can turn an absolute data set into a compositional data set by dividing each element of the sample vector by the total sum. (Figure 2 shows how the relationship between the samples (represented as points) changes when made compositional. Although the two groups appear clearly linearly separable in absolute space, the boundaries between groups become unclear in relative space. Meanwhile, the distances between samples become arbitrary.

When analyzing compositional data, it is sometimes possible to reclaim the discriminatory potential of relative data through transformation. For example, by setting all or some of the features relative to (i.e., divided by) a reference feature, one might discover that the resultant ratios can separate the groups ([66]). In fact, any separation revealed by such ratios can be analyzed by standard statistical techniques ([66]). This illustrates the concept behind the additive log-ratio (alr) transformation, achieved by taking the logarithm of each measurement within a composition (i.e., each sample vector containing relative measurements) as divided by a reference feature (i.e., *x_D_*) ([2]):

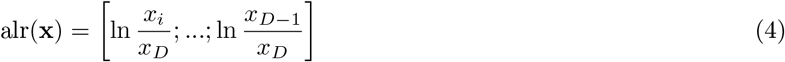

Instead of a specific reference feature, one could use an abstracted reference. In the case of the centered log-ratio (clr) transformation, the geometric mean of the composition (i.e., sample vector) is used in place of *x*_*D*_ ([2]). We use the notation *g*(x) to indicate the geometric mean of the sample vector, x. Note that because these transformations apply to each sample vector independently, the presence of an outlier sample does not alter the transformation of the other samples:

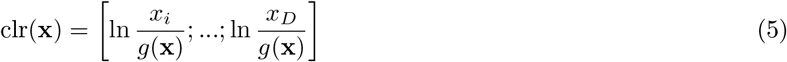

Likewise, other transformations exist that use the geometric mean of a feature subset as the reference. For example, the ALDEx2 package introduces the inter-quartile log-ratio (iqlr) transformation, which includes only features that fall within the inter-quartile range of total variance in the geometric mean calculation ([24]; [25]). Another, more complex, transformation, called the isometric log-ratio (ilr) transformation ([20]), also exists and is used in geological studies ([15]) and at least one analysis of RNA-Seq data ([67]). The ilr transforms the data with respect to an orthonormal coordinate system that is constructed from sequential binary partitions of features ([13]). Its default application to standard problems has been criticized by Aitchison on the basis that it lacks interpretability ([5]). Applications where the basis construction follows a microbiome phylogeny seem an interesting possibility ([74]).

### 4.3 The log-ratio “normalization”

In some instances, the log-ratio transformation is technically equivalent to a normalization. For example, let us consider the case where we know about our data the identity of a feature with a fixed abundance in absolute space across all samples. We could then use a log-ratio procedure to “sacrifice” this feature in order to “back-calculate” the absolute abundances. This is akin to using the alr transformation as a kind of normalization. However, because a single unchanged reference is rarely available or knowable (although synthetic RNA spike-ins may represent one way forward ([37])), we could try to approximate an unchanged reference from the data. For this, one might use the geometric mean of a feature subset, thereby using a clr (or iqlr) transformation as if it were a normalization.

Although log-ratio “normalizations” differ from log-ratio transformations only in the interpretation of their results, transformations alone are still useful even when they do not normalize the data. This is because they provide a way to move from the simplex into real space ([4]), rendering Euclidean distances meaningful. Importantly, clr- and ilr-transformed data impart four key properties to analyses: *scale invariance* (i.e., multiplying a composition by a constant k will not change the results), perturbation invariance (i.e., converting a composition between equivalent units will not change the results), *permutation invariance* (i.e., changing the order of the components within a composition will not change the results), and *sub-compositional dominance* (i.e., using a subset of a complete composition carries less information than using the whole) ([13]). Yet, the interpretation of transformation-based analyses remains complicated because the analyst must consider their results with respect to the chosen reference, or otherwise translate the results back into compositional terms.

### 4.4 Measures of distance

Euclidean distances do not make sense for compositional data ([4]). In contrast, the Aitchison distance does, providing a measure of distance between two *d*-dimensional compositions, x and **X** ([4]):

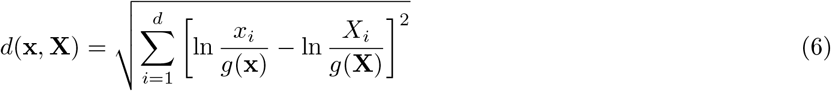

Although the Aitchison distance is simply the Euclidean distance between clr-transformed compositions, this distance (unlike Euclidean distance) has scale invariance, perturbation invariance, permutation invariance, and sub-compositional dominance. Few other distance measures satisfy all four of these properties, including none of the metrics routinely used in hierarchical clustering ([45]) (a routine part of RNA-Seq analysis). The property of sub-compositional dominance is especially important: even if the log-ratio transformation does not normalize the data, the addition of more sequence data will never make two samples appear *less* distant. This follows logically: as the amount of information available grows, the distance between samples should not shrink.

### 4.5 Measures of association

Like the Aitchison distance, there also exists a compositionally valid measure of association: the log-ratio variance (VLR) measures the agreement between two components (**a** and **b**) across two or more compositions. Specifically, it computes the variance of the logarithm of one component as divided by a second component. As such, a *D*-component data set contains *D*^2^ associations (albeit with symmetry). Unlike Aitchison distance, however, the VLR does not require a log-ratio transformation whatsoever; in fact, if using log-ratio transformed data, the reference denominators would cancel out. Note that, while distances occur between compositions (i.e., between samples), associations occur between components (i.e., between transcripts).

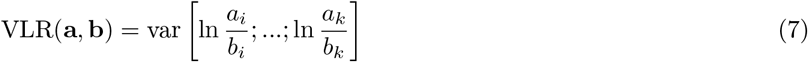

We can gain an intuition of the VLR by considering its formula. Recall that the relationship between components is one of relative importance: for the feature pair [**a**, **b**], the coordinates [2,4] and [4, 8] have equivalent meaning. Therefore, it follows that the features **a** and **b** are associated if 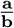 remains constant across all samples. Hence, we measure the variance of the (log-) ratios, such that VLR ranges from [0, inf] where 0 indicates a perfect association. Unfortunately, VLR lacks an intuitive scale, making non-zero values difficult to interpret ([43]).

Importantly, the VLR is *sub-compositionally coherent*: the removal of a third feature c would have no bearing on the variance of the (log-)ratio 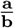. Yet, the VLR suffers from a key limitation: it is unscaled with respect to the variances of the log components ([43]). In other words, the magnitude of VLR depends partially on the variances of its constituent parts (i.e., var(**a**) and var(**b**)). This makes it difficult to compare VLR across pairs (e.g., comparing 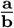 with 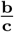) ([43]). Still, unlike correlation, the VLR does not produce spurious results for compositional data, and in fact, provides the same result for both relative data and the absolute counter-part, all without requiring normalization or transformation.

### 4.6 Principal Component Analysis

Just as there are problems regarding between-sample distances and between-feature correlations, it follows that Principal Component Analysis (PCA) should not get applied directly to compositional data. Instead, analysts could apply PCA to clr-transformed data (resulting in an additional centering of the rows after log-transformation) ([6]). However, analysts must take care when interpreting the resultant PCA: covariances and correlations between features now exist with respect to the geometric mean reference. As such, when plotting features as arrows in the new coordinate space, the angles between them (i.e., the correlations) will usually change when subsets of the data are analyzed. However, the distances between feature pairs (i.e., the links between the arrow heads) remain invariable with respect to sub-compositions: these correspond to their log-ratio variance ([6]). Meanwhile, the usual PCA plot (with samples as points in a new coordinate space) projects the distances between samples using the Aitchison distance (which has the desired property of sub-compositional dominance). In combining these into a joint visualization of features and samples, the resultant log-ratio biplot (i.e., the “relative variation biplot”) reveals associations between samples and features, and can also be used to infer power law relationships between features in an exploratory analysis ([6]). Such biplots are reminiscent of the visualizations obtained by Correspondence Analysis (CA). In fact, CA can indeed be used to approximate relative variation biplots provided the data are raised to a (small) power ([30]), the optimal size of which can be obtained by analyzing sub-compositional incoherence ([31]). Using CA with power transformation has the advantage that zeros in the data are handled naturally by the technique.

## 5 Compositional methods for sequence data

### 5.1 Methods for differential abundance

The ALDEx2 package, available for the R programming language, uses compositional data analysis principles to measure differential expression between two or more groups ([24]; [25]). Unlike conventional approaches to differential expression, ALDEx2 uses log-ratio transformation instead of effective library size normalization. The algorithm has five main parts. First, ALDEx2 uses the input data to create randomized instances based on the compositionally valid Dirichlet distribution ([24]; [25]). This renders the data free of zeros. Second, each of these so-called Monte Carlo (MC) instances undergoes log-ratio transformation, most usually clr or iqlr transformation ([24]; [25]). Third, conventional statistical tests (i.e., Welch's t and Wilcoxon tests for two groups; glm and Kruskal-Wallis for two or more groups) get applied to each MC instance to generate *p*-values (*p*) and Benjamini-Hochberg adjusted *p*-values (BH) for each transcript ([24]; [25]). Fourth, these p-values get averaged across all MC instances to yield expected *p*-values ([24]; [25]). Fifth, one considers any transcript with an expected BH < α as statistically significant ([24]; [25]).

Although popular among meta-genomics researchers for analyzing the differential abundance of operational taxonomic units (OTUs) (e.g., [48]; [70]), the ALDEx2 package has not received wide-spread adoption in the analysis of RNA-Seq data. In part, this may have to do with our observation that ALDEx2 requires a large number of samples. This requirement may stem from its use of non-parametric testing, as suggested by the reduced power of other non-parametric differential expression methods ([16]; [75]), for example NOISeq ([64]). However, competing software packages like limma ([62]) and edgeR ([56]) also benefit from moderated t-tests that “share information between genes” to reduce per-transcript variance estimates and increase statistical power.

Still, even in the setting of large sample sizes, ALDEx2 has one major limitation: its usefulness depends largely on interpreting the log-ratio transformation as a normalization. If the log-ratio transformation does not sufficiently approximate an unchanged reference, the statistical tests will yield results that are hard to interpret. Another tool developed for analyzing the differential abundance of OTUs suffers from a similar limitation: ANCOM ([44]) uses presumed invariant features to guide the log-ratio transformation. The tendency to interpret differential abundance results as if they were derived from log-ratio “normalizations” highlights the importance of pursuing numeric and experimental techniques that can establish an unchanged reference. It also highlights the benefit of seeking novel methods that do not require using log-ratio transformations as a kind of normalization.

### 5.2 Methods for association

The SparCC package, available for the R programming language, replaces Pearson's correlation coefficient with an estimation of correlation based on its relationship to the VLR (and other terms) ([28]). The algorithm works by iteratively calculating a “basis correlation” under the assumption that the majority of pairs do not correlate (i.e., a sparse network) ([28]). Another algorithm, SPIEC-EASI, makes the same assumption that the underlying network is sparse, but bases its method on the inverse covariance matrix of clr-transformed data ([38]).

The propr package ([53]), available for the R programming language, implements proportionality as introduced in ([43]) and expounded in ([22]). Proportionality provides an alternative measure of association that is valid for relative data. One could think of proportionality as a modification to the VLR that uses information about the variability of individual features (gained by a log-ratio transformation) to give the VLR scale. It can be defined for the *i*-th and *j*-th features (e.g., transcripts) of a log-ratio transformed data matrix, ã_*i*_ and ã_*j*_, and thus also depends on the reference used for transformation. Unlike SparCC and SPIEC-EASI, proportionality does not assume an underlying sparse network.

At least three measures of proportionality exist. The first, *ϕ*, ranges from [0,inf] with 0 indicating perfect proportionality ([43]):

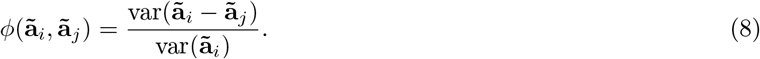

Its definition adjusts the VLR (in the numerator) by the variance of one of the log-ratio transformed features in that pair (in the denominator). The use of only one feature variance in the adjustment makes *ϕ* asymmetric (i.e., *ϕ*(ã_*i*_, ã_*j*_) ≠ *ϕ* (ã_*j*_, ã_*i*_)).

The second, *ϕ*_s_, also ranges from [0, inf] with 0 indicating perfect proportionality, but has a natural symmetry ([53]). Its definition adjusts the VLR by the variance of the log-product of the two features:

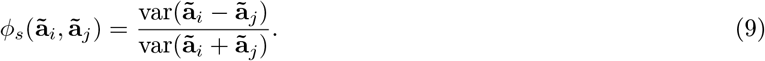

The third, *ρ*_*p*_, like correlation, takes on values from [-1,1], where a value of 1 indicates perfect proportionality ([22]). Its definition adjusts the VLR by the sum of the variances of the log-ratio transformed features in that pair (as subtracted from the value 1). Thus, *ρ*_*p*_ is symmetric.

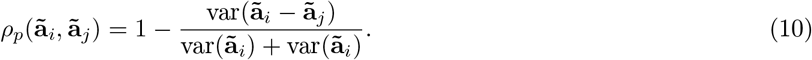

Note that *ρ*_*p*_ and ϕ_s_ are monotonic functions of one another (i.e., you can compute *ρ*_*p*_ directly from *ϕ_s_* and *vice versa*) (e.g., see ([22]) where *ϕ_s_* is called 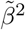). Unlike Pearson's correlation coefficient, proportionality coefficients tend not to produce spurious results ([53]). Instead, proportionality serves as a robust measure of association when analyzing relative data ([43]). Although proportionality gives VLR scale, it is limited in that its interpretation still depends partly on using transformation as a kind of normalization (i.e., for the calculation of individual feature variances) ([22]). Still, its interpretability, along with its observed resilience to spurious results, makes it a good choice for inferring co-expression from RNA-Seq data ([43]) or co-abundance from meta-genomics data ([10]).

## 6 Challenges to compositional analyses

### 6.1 Challenges unique to count compositions

Compositional data analysis, because it relies on log-transformations, does not work when the data contain zeros. Yet, count compositional data are notably prone to zeros, those of which could signify either that a component is absent from a sample or otherwise only present at a quantity below the detection limit ([14]). For NGS abundance data, the difference between a zero and a one might be stochastic. How best to handle zeros remains a topic of ongoing research. However, it is common to replace zeros with a number less than the detection limit ([14]). Other replacement strategies would include adding a fixed value to all components, replacing zeros with the value one, or omitting zero-laden components altogether. A more principled (yet computationally expensive) way of replacing zeros is the Dirichlet sampling procedure implemented in ALDEx2 (as described above). Note that the simple addition of a pseudo-count to all components does not preserve the ratios between them, which can be amended by modifying the non-zero components in a multiplicative way ([46]).

Moreover, while count compositional data carry relative information, they differ from true compositional data in that they contain integer values only. Restricting the data to integer space can introduce problems with an analysis because the sampling variation becomes more noticeable as the measurements approach zero ([53]). In other words, the difference between 1 and 2 counts is not exactly the same as the difference between 1,000 and 2,0000 counts ([53]). While it is not mathematically necessary to remove low counts, analysts should proceed carefully in their presence.

### 6.2 Challenges unique to sequencing data

In the second section, we discussed how between-sample biases render NGS abundances incomparable between samples, thus necessitating normalization or transformation. However, we did not address two important sources of within-sample biases for sequencing data. The first is read length bias, in which more reads map to longer transcripts ([63]). The second is GC content bias, in which more reads map to high GC regions ([19]). Such biases distort the ratios between features and are thus relevant to compositional analysis as well. Yet, because within-sample biases are usually assumed to have the same proportional impact across all samples, they are usually ignored ([63]). For the same reason, one might also ignore these biases when interpreting NGS abundance data as compositions (as long as we are only interested in between-sample effects). However, if a sample were to contain, for example, a polymorphic or epigenetic change which alters the size or GC content of a transcript, the compositional nature of sequencing data could cause a skew in the observed abundances for all other transcripts (for reasons suggested by (Figure 1). More work is needed to understand the extent to which within-sample biases impacts compositional data analysis in practice.

### 6.3 Limitations of transformation-based analysis

Formal transformation-based approaches often suffer from a lack of interpretability or otherwise get interpreted erroneously. For example, when using the centered log-ratio (clr) transformation, one may be tempted to interpret the transformed data as if they referred to single features (e.g., transcripts); however, the transformed data actually refer to the ratios of the transcripts to their geometric mean. As such, an analyst must interpret results with regard to their dependence on this mean. Moreover, because the geometric mean can change with the removal of features, the transformed data are incoherent with respect to sub-compositions.

When log-ratio transformations are used for scaled measures of association (i.e., proportionality), the resulting covariations depend on the implicitly chosen reference. Therefore, they will not give the same results for absolute and relative data (unless both data were transformed). The formal relationship of results when applying *ρ*_*p*_ with and without transformation is investigated elsewhere ([22]). Although lacking a natural scale, the log-ratio variance (VLR) has an advantage in that it provides identical results for both absolute and relative data, without requiring normalization or transformation.

### 6.4 The merits of ratio-based analysis

Aitchison's preferred summary of the covariance structure of a compositional data set was a matrix containing the log-ratio variances for all feature pairs (i.e., the variation matrix) ([2]). Although this matrix formally contains a lot of redundant information, an analyst who is familiar with the features might still find this kind of representation useful. Recently, the focus on ratios has been called the “pragmatic” approach to compositional data analysis ([32]), and offers some benefits. For one, transformation (i.e., the restriction to ratios with the same denominator) is not needed. Instead, the ratios can be dealt with directly as if they were unconstrained (i.e., absolute) data ([66]). Moreover, ratios may carry a clear meaning to the analyst interpreting them. Recently, Greenacre proposed a formal procedure to select a non-redundant subset of feature pairs that contains the entire variability of the data ([32]).

Such ratio-based analyses are also applicable to NGS abundance data. For example, Erb et al. proposed a method to identify the differential expression of gene ratios, a technique comprising part of what is termed differential proportionality analysis ([23]). When comparing gene ratios across two groups, this method selects ratios in which only a small portion of the total log-ratio variance (i.e., VLR) is explained by the sum of the within-group log-ratio variances ([23]). These selected gene ratios tend to show differences in the group means of those ratios, analogous to how genes selected by differential expression analysis show differences between their means ([23]). Reinforcing the analogy further, Erb et al. have shown how it is possible to use the limma package to apply an empirical Bayes model with underlying count-based precision weights ([62]; [39]) to gene ratios, thus quantifying “second order” expression effects while still avoiding normalization ([23]).

In addition to measuring differences in the means of gene ratios between groups, ratio-based methods (such as those used in differential proportionality analysis) can also help identify differences in the coordination of gene pairs. Such “differential coordination analysis” would otherwise depend on correlation ([76]), and therefore fall susceptible to spurious results. Instead, we can harness the advantages of the VLR to define a sub-compositionally coherent measure that tests for changes in the magnitude (i.e., slope of association) or strength (i.e., coefficient of association) of co-regulated gene pairs. Moreover, ratio-based analyses could work as normalization-free feature selection methods for data science applications (such as clustering and classification). Such techniques would especially suit large data sets aggregated from multiple sequencing centers, platforms, or modalities, where heterogeneity and batch effects are not easily normalized.

## 7 Summary

All NGS abundance data are compositional because sequencers sample only a portion of the total input material. However, RNA-Seq data might have compositional properties regardless owing to constraints on the cellular capacity for mRNA production. Whatever the reason, compositional data cannot undergo conventional analysis directly, at least without prior normalization or transformation. Otherwise, measures of differential expression, correlation, distance, and principal components become unreliable.

In the analysis of RNA-Seq data, effective library size normalization is used to recast the data in absolute terms prior to analysis. However, successful normalization requires meeting certain (often untestable) assumptions. Alternatively, log-ratio transformations provide a way to interrogate the data using familiar methods, but analysts must interpret their results with respect to the chosen reference. Sometimes, log-ratio transformations can be used to normalize the data, but this requires an approximation of an unchanged reference. Instead, shifting focus to the analysis of ratios yields methods that avoid normalization and transformation entirely. These ratio-based methods may represent an important future direction in the compositional analysis of relative NGS abundance data.

